# Evaluating brain structure traits as endophenotypes using polygenicity and discoverability

**DOI:** 10.1101/2020.07.17.208843

**Authors:** Nana Matoba, Michael I. Love, Jason L. Stein

## Abstract

Human brain structure traits have been hypothesized to be broad endophenotypes for neuropsychiatric disorders, implying that brain structure traits are comparatively ‘closer to the underlying biology’. Genome-wide association studies from large sample sizes allow for the comparison of common variant genetic architectures between traits to test the evidence supporting this claim. Endophenotypes, compared to neuropsychiatric disorders, are hypothesized to have less polygenicity, with greater effect size of each susceptible SNP, requiring smaller sample sizes to discover them. Here, we compare polygenicity and discoverability of brain structure traits, neuropsychiatric disorders, and other traits (89 in total) to directly test this hypothesis. We found reduced polygenicity (FDR = 0.01) and increased discoverability of cortical brain structure traits, as compared to neuropsychiatric disorders (FDR = 3.68×10^−9^). We predict that ~8M samples will be required to explain the full heritability of cortical surface area by genome-wide significant SNPs, whereas sample sizes over 20M will be required to explain the full heritability of major depressive disorder. In conclusion, we find reduced polygenicity and increased discoverability of cortical structure compared to neuropsychiatric disorders, which is consistent with brain structure satisfying the higher power criterion of endophenotypes.

## Introduction

Human brain structure traits have been posited to be broad endophenotypes for neuropsychiatric disorders [Almasy and Blangero, 2001; Bigos and Weinberger, 2010; Flint and Munafò, 2007; Meyer-Lindenberg and Weinberger, 2006]. Endophenotypes have two attractive properties for genetic search [Le and Stein, 2019]: First, *higher power*, because precisely measured endophenotypes are ‘closer to the underlying biology’ than heterogeneous, clinically defined disorders, smaller sample sizes are needed to detect endophenotype effects. Second, *mechanistic insight*, because those variants associated with an endophenotype also influence risk for neuropsychiatric disorders, endophenotype associations are informative about the mechanisms leading to risk for neuropsychiatric disorders. Genome-wide association studies (GWAS) have identified common genetic variants associated with many traits, including brain structure [Adams et al., 2016; Elliott et al., 2018; Grasby et al., 2020; Hibar et al., 2015; Hibar et al., 2017; Satizabal et al., 2019; Stein et al., 2012; Zhao et al., 2020] and risk for neuropsychiatric disorders [Demontis et al., 2019; Howard et al., 2019; Matoba et al., 2020; Pardiñas et al., 2018; Stahl et al., 2019]. GWAS results from large sample sizes allow for the comparison of common variant genetic architectures between traits [Watanabe et al., 2019] and the direct evaluation of these endophenotype properties.

Genetic architecture can be summarized by several parameters [Holland et al., 2020; Zhang et al., 2018]: (1) *heritability (h^2^)*: the overall amount of trait variance explained by genetics, (2) *polygenicity* (*π_c_*): the proportion of susceptibility SNPs (sSNPs), LD-independent loci associated with a trait that are not necessarily genome-wide significant, relative to the total number of LD-independent SNPs in the genome (M) and (3) *discoverability* (σ): the distribution of effect sizes of sSNPs on a trait. Higher polygenicity of a trait indicates more sSNPs that are associated with that trait (**Figure 1**). Higher polygenicity is generally associated with lower effect size of each sSNP, requiring higher sample sizes to discover them [Watanabe et al., 2019]. Endophenotypes, compared to neuropsychiatric disorders, are hypothesized to have less polygenicity, with greater effect size of each sSNP, requiring lower sample sizes to discover them.

**Figure 1:**
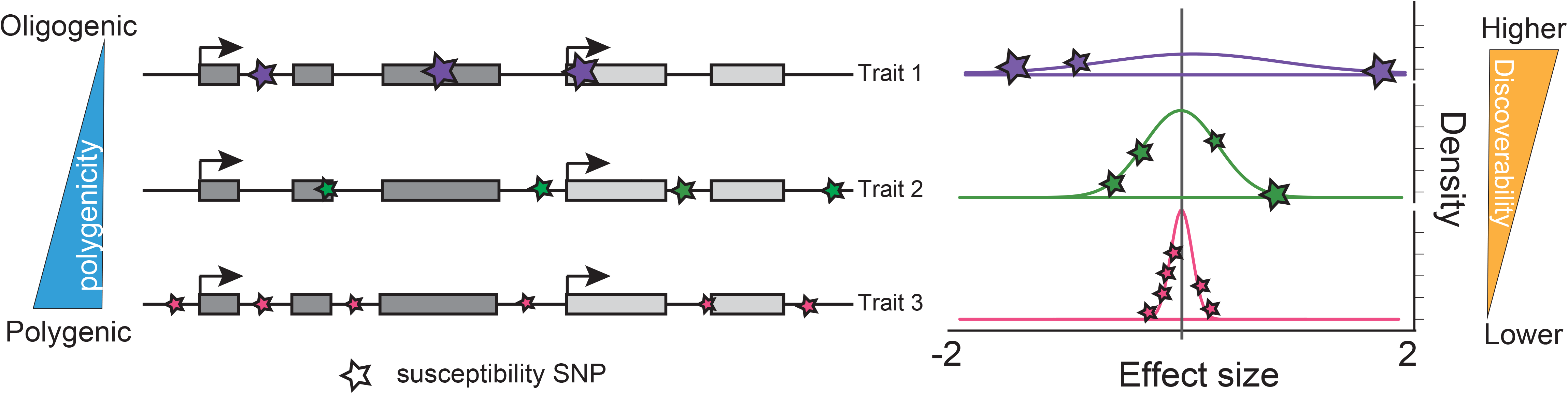
Illustration of polygenicity and discoverability. For traits with higher polygenicity, more variants (LD independent susceptibility SNPs indicated as stars, left) are associated with the trait, each at lower effect size which leads to lower discoverability (right).

Here, we directly tested whether brain structure traits satisfy the higher power property of endophenotypes using summary statistics from existing GWAS. We applied GENESIS [Zhang et al., 2018], a mixture model algorithm that clusters LD independent SNPs into either null SNPs, those that have no detectable effect on a trait, or sSNPs, those that have a detectable influence without requiring genome-wide significance. We found reduced polygenicity and increased discoverability of cortical surface area traits as compared to both cortical thickness and subcortical volumes. Additionally, we found reduced polygenicity and increased discoverability of cortical brain structure traits, as compared to neuropsychiatric disorders and anthropometric traits. We therefore project that, as additional GWAS are completed in the future, studies will find more explained heritability of cortical structure traits compared to equivalently-sized studies of neuropsychiatric disorders. These findings support cortical brain structure traits as satisfying the higher power criteria of an endophenotype.

## Methods

### GWAS summary statistics

We obtained GWAS summary statistics for 89 complex traits and disorders. Summary statistics for brain structure traits including cortical surface area (n = 35), thickness (n = 35) [Grasby et al., 2020] and subcortical volumes (n = 7) [Hibar et al., 2017; Satizabal et al., 2019] were obtained from the Enhancing NeuroImaging Genetics through Meta Analysis (ENIGMA) consortium (http://enigma.ini.usc.edu/research/download-enigma-gwas-results/). From the Psychiatric Genomics Consortium (PGC) (https://www.med.unc.edu/pgc/download-results/), summary statistics for three psychiatric disorders (schizophrenia [Ripke et al., 2013; Schizophrenia Working Group of the Psychiatric Genomics Consortium, 2014], bipolar disorder [Stahl et al., 2019], depression [Howard et al., 2019; Wray et al., 2018] (including major depression (MDD) and broad depression, excluding 23andMe participants)) were obtained. We additionally downloaded summary statistics for schizophrenia [Pardiñas et al., 2018] from https://walters.psycm.cf.ac.uk/. Summary statistics for Attention deficit/hyperactivity disorder (ADHD) (European population) [Demontis et al., 2019] were obtained from the Integrative Psychiatric Research (iPSYCH) website (https://ipsych.dk/en/research/downloads/). Summary statistics for Autism Spectrum Disorder (ASD) were generated in our previous study [Matoba et al., 2020]. Summary statistics for addiction (cigarettes per day and drinks per week) [Liu et al., 2019] were downloaded from the GWAS & Sequencing Consortium of Alcohol and Nicotine use (GSCAN) (https://conservancy.umn.edu/handle/11299/201564). Summary statistics for cognitive function (intelligence [Savage et al., 2018] and reaction time [Watanabe et al., 2019]) were obtained from https://ctg.cncr.nl/software/summary_statistics and https://atlas.ctglab.nl/ukb2_sumstats/f.20023.0.0_res.EUR.sumstats.MACfilt.txt.gz, respectively. Summary statistics for anthropometric measurements (height and BMI) [Yengo et al., 2018] were obtained from the Genetic Investigation of ANthropometric Traits (GIANT) consortium (https://portals.broadinstitute.org/collaboration/giant/index.php/GIANT_consortium_data_files#BMI_and_Height_GIANT_and_UK_BioBank_Meta-analysis_Summary_Statistics) Further information is summarized in **Supplementary Table 1**.

### Data preparation for GENESIS

The proportion of sSNPs for 89 traits and diseases and their effect size distributions were estimated using GENetic Effect-size distribution Inference from Summary-level data (GENESIS; v1.0) (https://github.com/yandorazhang/GENESIS) [Zhang et al., 2018]. GENESIS is a tool that distinguishes sSNPs from null SNPs using a mixture model of effect sizes from GWAS summary statistics in order to estimate parameters describing the genetic architecture of a trait. As the software requires rsID, Z score, and effective sample size of GWAS study as inputs, we calculated Z scores using effect sizes (beta or log(OR)) and standard errors (se). If the downloaded summary statistics did not provide the sample numbers for individual SNPs, we used the total number of enrolled participants (# of cases and controls for case-control studies). For case-control studies, effective sample sizes were further estimated by 4/(1/cases + 1/controls) [Willer et al., 2010]. Using the SNP QC function (*preprocessing*) implemented in GENESIS, SNPs with a low effective sample size (< 0.67 * 90^th^ percentile of sample size), or very large effect size (*Z^2^ > 80*) were removed. This function also removed SNPs within the major histocompatibility complex (MHC) region. Only those SNPs in HapMap3 [The International HapMap 3 Consortium, 2010] with MAF ≥ 0.05 in European population from the 1000 Genome project Phase 3 (1KG) [The 1000 Genomes Project Consortium, 2015] were retained. We used pre-computed LD-scores, which were also estimated from common SNPs in HapMap 3 using LD from 1KG European population as described in the original GENESIS paper [Zhang et al., 2018].

### Model selection

We ran the *genesis*() function with default options (LDcutoff (*r^2^*) = 0.1, LDwindow = 1 Mb, M = 1,070,777 total number of reference SNPs). GENESIS implements two models (the two-component model, M2; and the three-component model, M3), which assumes that the distribution of effects for non-null SNPs follows either a single normal distribution or mixture of two normal distributions (allowing two distinct sSNP groups based on effect size). Variance parameters for the M3 model were estimated using output from the M2 model as recommended in the GENESIS documentation. To select the best fit model, we used the modified Baysesian information criterion (BIC), also implemented in GENESIS [Zhang et al., 2018] as well as the ratio of variance estimates from the M3 model. We used M2 if the ratio of two variance estimates 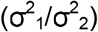 from M3 was less than 5 or if the BIC for M2 was less than M3. (**Supplementary Figure 1-5, Supplementary Table 2**). QQ plots were generated to evaluate goodness of model fit by comparing p-values from the GWAS summary statistics with the fit model estimates. Expected p-values from the fit models and 80% confidence intervals were internally generated in *genesis*().

### Estimation of polygenicity and effect-size distributions

After selecting the best model, we then estimated the parameters of genetic architecture: polygenicity and discoverability. The mixture model provides the proportion of non-null SNPs (sSNPs) for each trait, which is the polygenicity (*π_c_*). The total number of sSNPs was estimated by

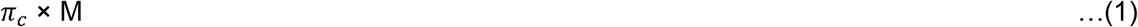

where *π_c_* is the proportion of sSNPs obtained by *genesis*() and M is the number of SNPs in the reference panel (M = 1,070,777). And the number of sSNPs in the cluster with larger variance component for M3 was estimated by multiplying the proportion of sSNPs in that cluster (**Supplementary Table 3**). 95% CIs for number of sSNPs were also calculated by adding and subtracting 1.96 times the standard error for *π_c_* and plugging in these interval endpoints into formula (1). We note the standard error of *π_c_* for cuneus thickness was not able to be estimated by genesis, so we could not estimate the 95% CI of *π_c_* and number of sSNPs for this trait.

In order to compare effect size distributions across traits (discoverability), regardless of whether the traits were modeled with M2 or M3, we selected one quantity from the distribution: the 50th percentile of ranked sSNPs absolute effect size. In other words, the predicted effect size of an sSNP where half of all sSNPs have larger effect size in absolute value. In order to estimate this quantity, we used the distribution and quantile functions in R-3.5.0. For phenotypes with M2, we used

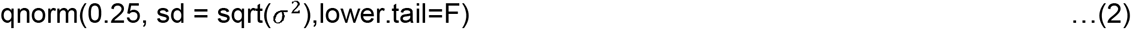

where *σ*^2^ is variance estimated by GENESIS

For phenotypes with M3, we found the smallest value “x” via grid search such that

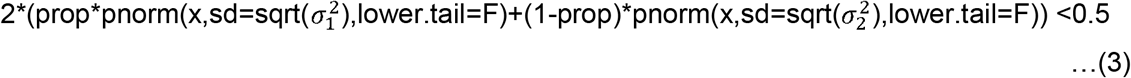

where prop is the proportion of sSNPs in cluster 1 with larger effect sizes, 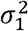 is the variance in cluster 1, 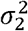 is variance in cluster 2 with smaller effect sizes, and s = seq(0,0.02, length=200). 95% confidence intervals (CI) for each parameter were calculated by adding and subtracting 1.96 times the standard error for each parameter (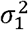, 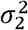, prop) output from GENESIS, and plugging in these interval endpoints into the formula above ((2) or (3) based on best fit model). For some traits, the range of CIs were outside possible values, so we limited them as follows: (1) if the lower bound of 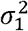 or 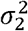 was < 0, we set its value to 0, (2) if the lower bound of proportion of sSNPs in cluster 1 was < 0, we set its value to 0 (meaning that the sSNPs were considered to belong to cluster 2) and (3) if the upper bound of prop was > 1, we set its value to 1.

### Prediction of sample sizes needed to attain complete heritability

We estimated the predicted heritability explained by genome-wide significance (GWS) SNPs (GVpercentage) with a given sample size from 50,000 to 200,000,000 (interval = 50,000) by applying the *projection*() function in GENESIS. We defined the sample size needed to achieve complete heritability as the sample size required for the GVpercentage to pass 99%. We computed this prediction with the best fit model for each phenotype. If the GVpercentage did not pass 99% at a sample size of 200,000,000, we showed the GVpercentage achieved at that sample size.

### Comparison of estimates across categories

The implementation of GENESIS does not have functions that directly compare the polygenicity or discoverability across traits or groups of traits. In order to compare these values across traits, it is necessary to take into account the standard error of each parameter estimate. After standard errors were calculated, the heterogeneity (*I^2^* statistic) between groups of traits was calculated based on a fixed effect model implemented in *metagen*() function in Meta package (v4.12-0) [Schwarzer et al., 2015], and specifying the argument byvar=group. Because the standard error of *π_c_* for cuneus thickness was not able to be estimated by GENESIS (see above in estimation of polygenicity and effect-size distributions section), we excluded cuneus thickness and cuneus surface area to avoid potential biases in comparisons (**Figure 2**). The FDR-adjusted p-values [Benjamini and Hochberg, 1995] (FDR < 0.05) of heterogeneity across six pairs of trait groups was used to determine the significance. The outputs from metagen were further used to generate a forest plot.

**Figure 2:**
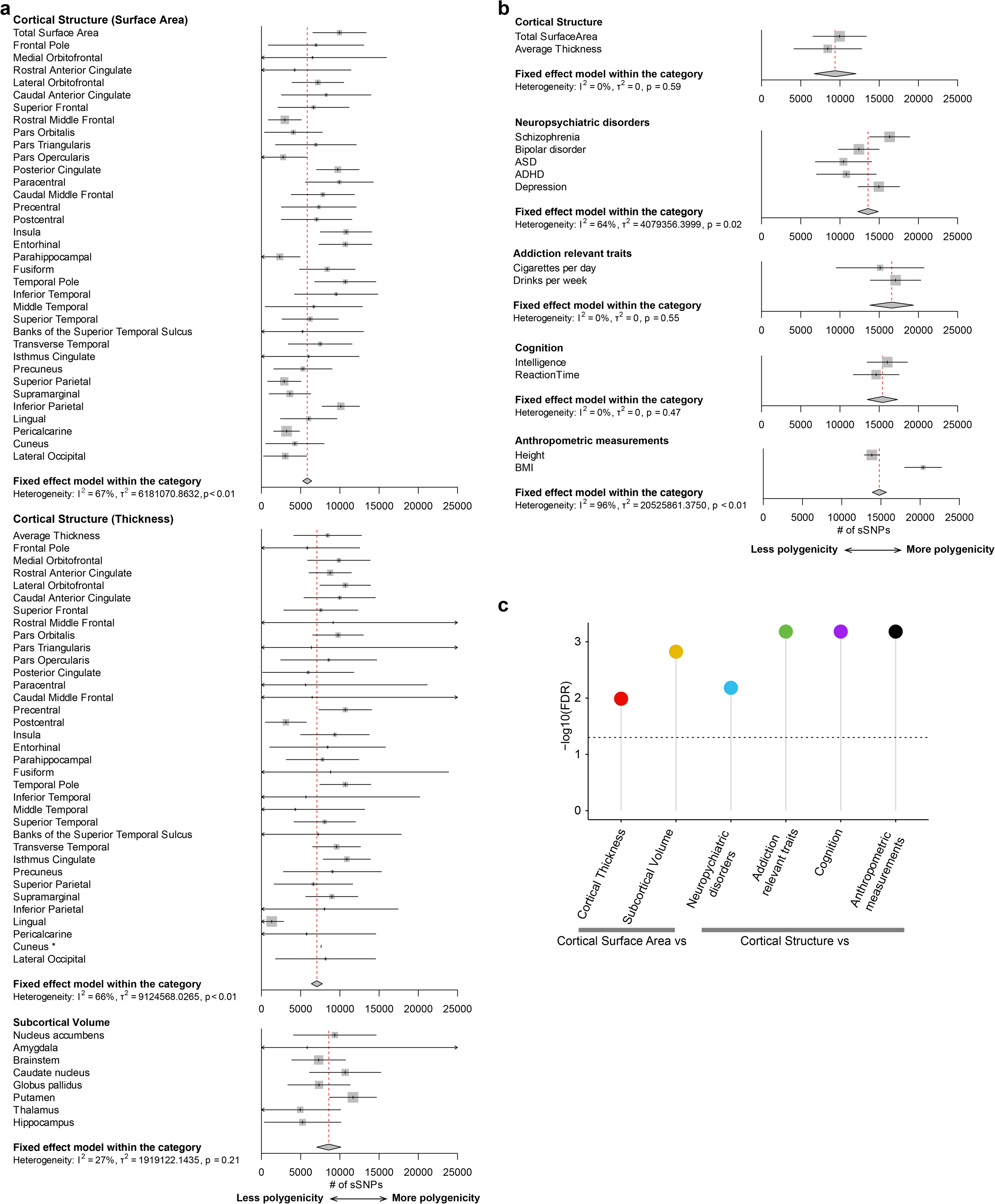
Estimates of polygenicity across multiple complex brain-relevant traits. The predicted number of sSNPs shows **(a)** decreased polygenicity for cortical surface area compared to cortical thickness or subcortical volumes, and **(b)** decreased polygenicity for global cortical traits compared to neuropsychiatric disorders, addiction traits, cognition, and anthropometric measurements. **(c)** The significance after FDR correction between categories, calculated via a heterogeneity test. The horizontal line indicates −log_10_(FDR = 0.05). Because standard error for polygenicity (*π_c_*) of the cuneus thickness was not able to be estimated, we excluded this region from the heterogeneity tests for cortical thickness. To avoid bias, we also excluded cuneus surface area when comparing cortical surface area and cortical thickness.

## Results

We first obtained GWAS summary statistics of various traits including cortical and subcortical brain structure [Grasby et al., 2020; Hibar et al., 2017; Satizabal et al., 2019]), neuropsychiatric disorders and cognitive phenotypes [Demontis et al., 2019; Howard et al., 2019; Liu et al., 2019; Matoba et al., 2020; Pardiñas et al., 2018; Ripke et al., 2013; Savage et al., 2018; Schizophrenia Working Group of the Psychiatric Genomics Consortium, 2014; Stahl et al., 2019; Watanabe et al., 2019; Wray et al., 2018], and anthropometric measurements [Yengo et al., 2018] (**Supplementary Table 1)**. All of the GWASs were performed in European ancestries. Effective sample size ranged from 29,235 individuals for brain stem volume to 795,640 individuals for body mass index (BMI). In order to quantify parameters of genetic architecture for each of these traits, we applied GENetic Effect-size distribution Inference from Summary-level data (GENESIS) [Zhang et al., 2018] to those GWAS summary statistics (**Supplementary Table 1**). Using GENESIS, for each trait, we estimated several parameters describing the genetic architecture of the traits: (1) polygenicity (*π_c_*) and the total number of sSNPs as (*π_c_* × M), (2) discoverability quantified as the variance of effect sizes for non-null sSNPs (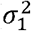 and 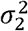), and (3) predictions of heritability explained by genome-wide significance (*P* < 5.0×10^−8^; GWS) sSNPs in future sample sizes. We then compared these genetic architecture parameters across traits.

### Model Selection

We first identified the best fit model comprising either one set of null SNPs and one set of sSNPs (M2) or one set of null SNPs and two sets of sSNPs at different levels of effect size (M3) for each trait (**Supplementary Table 2, 3**). Among 89 traits, the M3 model best fit 39 traits (44.9%). The thickness and surface area of the brain cortex (n = 35 traits each) showed somewhat different proportions of the best fit model (i.e. 65.7% of surface area GWAS best fit M2, while 60.0% of thickness GWAS best fit M3), though this difference was not significant (Fisher’s exact test; *p* = 0.055) (**Supplementary Figure 1, 2**, **Supplementary Table 2**). To evaluate goodness of model fit to the observed data, we generated Q-Q plots which allows visual assessment of whether the expected p-values from the model correspond to the empirically observed p-values from GWAS summary statistics (**Supplementary Figure 1-5**). Generally, we observed strong goodness of fit for the best fit model, where the observed p-values corresponded to the model p-values. However, for some traits fit to the M3 model (e.g. surface area of lateral orbitofrontal area and thickness of caudal middle frontal area), there are a number of outlier sSNPs (P < 10^−10^) implying that these traits could be fit to more complex models [Zhang et al., 2018].

### Comparing polygenicity across complex brain-relevant traits

We compared the number of susceptibility SNPs (sSNPs), a measure of polygenicity, between groups of related traits. We found that global surface area had *π_c_* = 0.9% with 9,949 sSNPs (95% CIs: 6,552 to 13,346) (**Figure 2a**). Current GWAS results have detected only 20 genome-wide significant loci [Grasby et al., 2020], but these results show that many more significant loci are expected to be associated with global surface area as sample sizes grow. We noted that there was heterogeneity in terms of polygenicity across different cortical regions with insula having the highest polygenicity (10,791 sSNPs, 95% CIs 7,510 to 14,072) and parahippocampal gyrus having the least polygenicity (2,365 sSNPs, 95% CIs 0 to 5,989). There was significant heterogeneity observed across the 35 cortical surface area traits, indicating regional genetic architecture varies even for the same measure (surface area) across cortical regions (*I^2^* = 67%; p < 0.01). We then compared polygenicity from the 34 cortical surface area traits to the 34 matched cortical thickness traits (excluding cuneus, see Methods). The predicted number of sSNPs for cortical surface area was significantly smaller than cortical thickness (FDR = 0.01; **Figure 2a, c**), indicating that cortical surface area has reduced polygenicity as compared to thickness. Similarly, subcortical volumes also have reduced polygenicity relative to cortical surface area, indicating that cortical surface area traits, as a group, are the least polygenic among these tested brain structure traits.

Next, we tested differences in polygenicity for global cortical structure traits (global thickness and surface area) compared to brain-relevant and anthropometric traits. Consistent with previous findings [Zhang et al., 2018], these neuropsychiatric disorders, addiction relevant traits, cognition, and anthropometric traits all had high levels of polygenicity (**Figure 2b**). Interestingly, we found significantly reduced polygenicity for cortical structure traits as compared to neuropsychiatric disorders (FDR = 0.007), addiction relevant traits (FDR = 0.0007), cognition (FDR = 0.0007) and anthropometric measurements (FDR = 0.0007) (**Figure 2b-c**). Overall these results are consistent with predictions of cortical structure as satisfying the higher power criterion of an endophenotype.

### Comparing discoverability across complex brain-relevant traits

We next examined the effect size distribution across the same traits. In **Figure 3a**, we plotted the estimated effect size distribution of global cortical structure phenotypes, neuropsychiatric disorders, cognition, addiction relevant traits, and brain relevant traits using the best fit model. As expected based on the polygenicity results, we observed increased discoverability, wider effect size distribution and larger value of the a parameter(s), of cortical structure traits as compared to others. However, this visual comparison does not include confidence intervals of discoverability estimates, limiting statistical inference across traits. To identify one quantity from the effect size distribution, including confidence intervals, that can be compared across traits potentially modeled with different mixture distributions, we used the absolute value of the effect size of the 50th percentile of ranked sSNPs (**Figure 3b**). Consistent with the polygenicity results above, we observed increased discoverability of effect sizes in cortical surface area compared to cortical thickness (FDR = 5.66×10^−5^). However, we did not observe statistically different effect sizes in cortical surface area compared to subcortical volumes (FDR = 0.05) (**Figure 3d**, **Supplementary Figure 6**).

**Figure 3:**
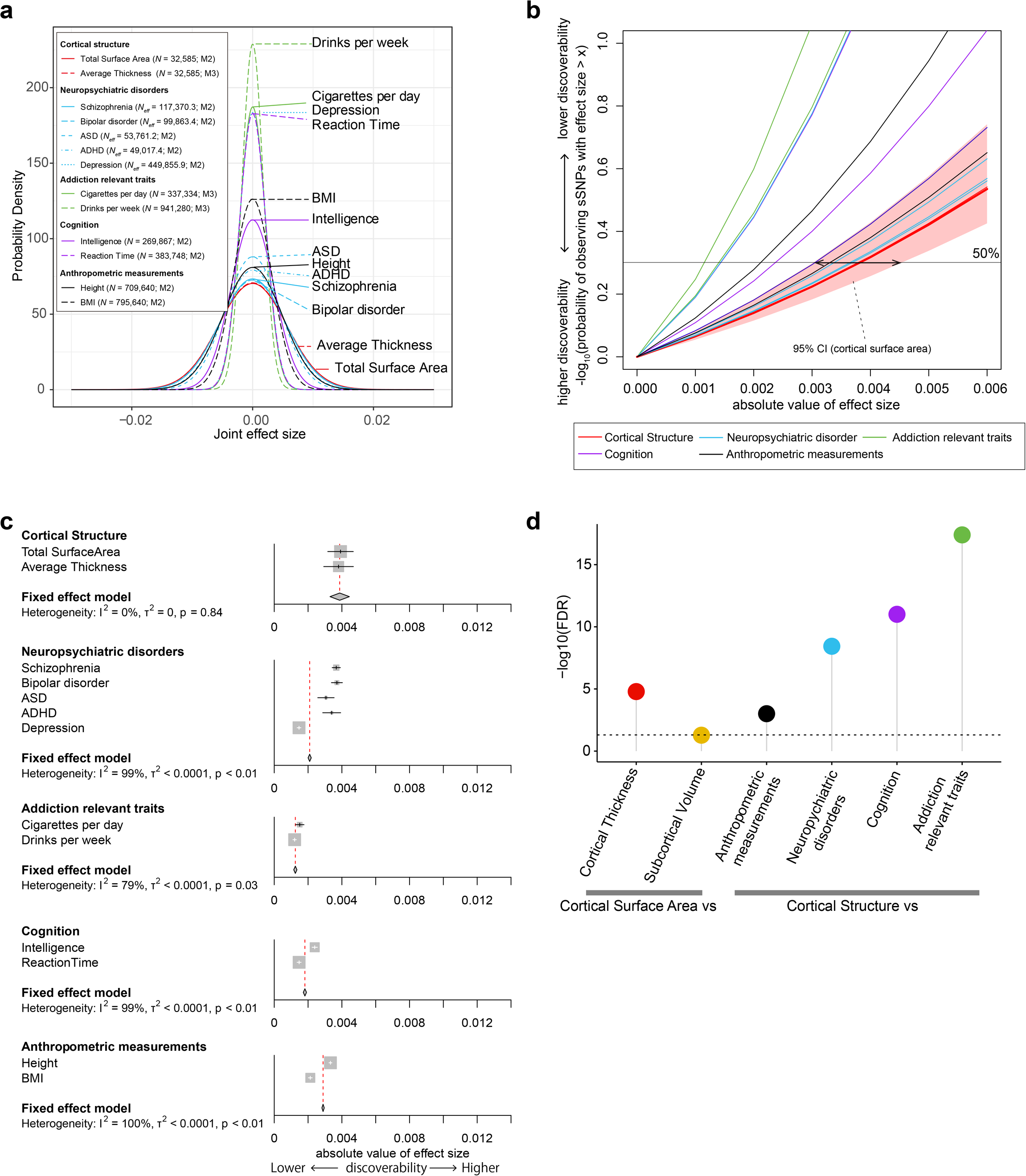
Estimates of discoverability across multiple complex brain-relevant traits. The effect size distributions across traits suggest increased effect sizes in cortical structure compared to complex traits including four neuropsychiatric disorders, two addiction relevant traits, two cognitive traits, and two anthropometric measurements (**a**-**d**). Joint effect sizes are an approximation of Pearson’s correlation coefficient between sSNPs and phenotype. (**a**) M2/M3 indicates the best fit model for the traits. The effect size distribution (variance) shows increased absolute effect size for global cortical traits compared to neuropsychiatric disorders, addiction traits, cognition, and anthropometric measurements. **(b)** The ranked absolute effect size at the 50th percentile of observed sSNPs were compared. The red horizontal line indicates the 50% probability. **(c)** A comparison of the absolute effect size at the 50th percentile across traits. Note that phenotypes annotated with a * require caution in interpretation because the lower limit of the 95% CI of proportion of sSNPs in cluster 1 (the larger variance component) was estimated to be a negative value and we limited it to 0 for these phenotypes (all sSNPs were considered in cluster 2 (smaller variance component) in this case). The significance between categories under FDR correction, calculated via a heterogeneity test, is displayed in **(d)**. The horizontal line indicates −log_10_(FDR = 0.05). See also **Supplementary Figure 6** for comparison across cortical/subcortical regions.

We also statistically compared the effect size distribution of genetic variants across brain-relevant traits including neuropsychiatric disorders, addiction related traits, cognitive function, and anthropometric measurements (**Figure 3c, d**). The heterogeneity tests indicated significantly increased effect sizes in cortical structure compared to those complex traits (vs neuropsychiatric disorders: FDR = 3.68 × 10^−9^; addiction relevant traits: FDR = 4.03 × 10^−18^; cognition: FDR = 9.88×10^−12^; anthropometric measurements: FDR = 9.80×10^−4^). There was also heterogeneity within each category, and the difference in observed significance between cortical structure and neuropsychiatric disorders came mainly from depression, insofar as the significance decreased when removing depression from the group of traits (FDR w/o depression = 0.33) (**Figure 3c, d**). In summary, we found evidence that sSNPs for brain structure have stronger effect size compared to neuropsychiatric disorders, largely driven by the low effect sizes observed in depression. We also observed that brain structure traits have stronger effect sizes when compared to cognition or addiction relevant traits.

### Correlation between polygenicity, discoverability and heritability

Previous studies have shown that increasing polygenicity is associated with decreased discoverability [Watanabe et al., 2019]. To test if this same relationship was observed among the traits tested here, we correlated these measures. We observed an inverse relationship between estimated polygenicity and estimated discoverability (Peason’s correlation coefficient (r) = −0.65, *p* = 4.43×10^−7^; **Supplementary Figure 8a**), though such a large negative correlation between polygenicity and discoverability was expected given the covariance of estimated parameters output by GENESIS (**Supplementary Figure 8b**). Moreover, discoverability was also correlated to heritability of brain-related phenotypes (r = 0.56, *p* = 1.57×10^−8^; **Supplementary Figure 8c**). Interestingly, we found no evidence that polygenicity and heritability are correlated (r = 0.067, *p* = 0.53; **Supplementary Figure 8c**) indicating that heritability might be independent of polygenicity but be affected by the magnitude of effect size of sSNPs.

### Sample sizes needed to explain full heritability

Next, we predicted the number of subjects needed in future GWAS to identify all of the common variant loci (sSNPs) associated with a trait. In other words, we estimated the sample size needed to achieve 99% heritability explained by genome-wide significant SNPs. (**Figure 4, Supplementary Table 4**). We predict that at least 8 million individuals will be required to explain the full heritability of global cortical surface area and 8.65 million for cortical thickness. Notably, while less than 20 million individuals will be needed to explain the full heritability for the majority of regional surface area traits, about half of regional cortical thickness will not achieve full heritability even at that large sample size. As expected, larger sample sizes will be needed to explain the full heritability for phenotypes that showed lower discoverability or increased polygenicity such as depression or drinks per day.

**Figure 4:**
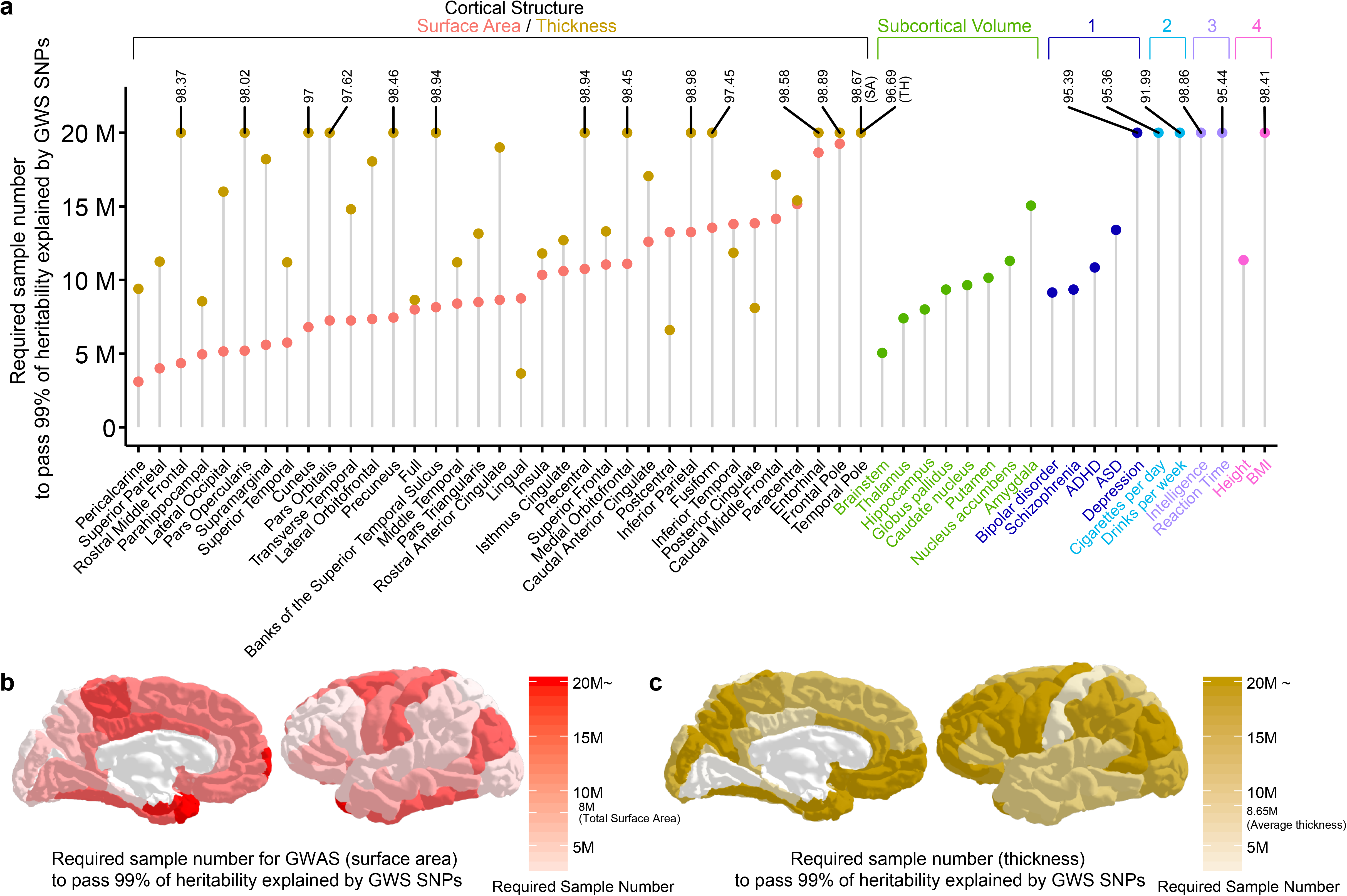
Predicted sample sizes required to explain the full heritability of traits. Predicted required sample size (up to 20M) for brain related phenotypes. **(a)** The y axis shows the predicted sample size to explain 99% of heritability explained by GWS sSNPs. Phenotypes that did not reach 99% were labeled with the predicted percentage of heritability explained at a sample size of 20M. 1. neuropsychiatric disorders, 2. addiction relative traits, 3. Cognition, 4. anthropometric measurements. Regions colored by sample sizes needed to attain full heritability for **(b)** cortical surface area or **(c)** thickness.

## Discussion

Here, we directly tested one proposed property of brain structure traits as endophenotypes: higher power of genetic discovery. We evaluated this by estimating the polygenicity and discoverability (effect size distribution) across multiple cortical and subcortical brain structure traits and compared these to the same measures from neuropsychiatric disorders, cognitive, addiction relevant, and brain related traits. We found that cortical structure traits have reduced polygenicity and increased discoverability compared to neuropsychiatric disorders. This is consistent with brain structure satisfying the higher power criterion of endophenotypes.

Our results have both practical and theoretical implications. Practically, brain structure traits have higher power than neuropsychiatric disorders so smaller sample sizes will be required to achieve equivalent gains in genetic discovery. The costs of phenotype acquisition for brain MRI are still high, but nevertheless large biobanks and integration of genotype data with electronic medical records are now developing that will allow GWAS of brain structure in sample sizes of hundreds of thousands in the near future [Bowton et al., 2014; Elliott et al., 2018]. Theoretically, we hypothesize that polygenicity and discoverability are related to the number of causal mechanisms that impact a trait. For example, genetic variants associated with molecular traits like chromatin accessibility have very high effect sizes and low polygenicity [Liang et al., 2020] - a single variant may explain most of the heritability of an accessible region. This is likely because there are a limited number of mechanisms by which genetic variation influences accessibility, namely the ability of DNA binding proteins (like transcription factors) to bind to the genome. Genetic variants associated with gene expression also have high effect sizes, but lower than caQTLs [Liang et al., 2020]. This is likely because multiple mechanisms can influence gene expression including transcription factor binding, miRNA expression levels, methylation, and RNA degradation [Li et al., 2016]. Extending this logic to complex traits like brain structure, we would predict that fewer mechanisms influence cortical surface area compared to neuropsychiatric disorders. For example, we previously have described evidence to support that cortical surface area is influenced by proliferation of neural progenitor cells present in fetal development [Grasby et al., 2020; de la Torre-Ubieta et al., 2018]. Whereas, for depressive disorder, which has some shared genetic basis with cortical surface area, we would hypothesize that multiple mechanisms including multiple cell-types (progenitors and mature neurons), multiple cellular processes (proliferation and neuronal firing), in multiple developmental and tissue contexts all create risk for the complicated disorder.

We note that our study was only designed to address the first criterion of an endophenotype, higher power. In order to gain mechanistic insight into the basis of neuropsychiatric disorders using brain structural traits, there must also be both a genetic correlation and evidence of mediation between the brain structural trait and risk for a neuropsychiatric disorder [Kendler and Neale, 2010; Le and Stein, 2019]. Significant genetic correlations have been demonstrated between ADHD, major depressive disorder, and brain structure traits [Grasby et al., 2020; Klein et al., 2019; Satizabal et al., 2019]. Notably though, no significant genetic correlations have yet been observed between brain structural traits and schizophrenia [Franke et al., 2016; Grasby et al., 2020], so it is unlikely that the brain structural traits explored in this study will provide mechanistic insight into the basis of schizophrenia.

We should interpret our results in light of some limitations. First, earlier studies have shown that estimates of effect sizes are likely to be biased upwards in smaller sample sizes (winner’s curse) [Kraft, 2008; O’Sullivan and Ioannidis, 2020; Xiao and Boehnke, 2009]. To examine the possibility that smaller sample sizes in brain structure traits may have inflated discoverability estimates, we compared discoverability estimates from historical schizophrenia GWAS with sample sizes ranging from N_eff_=31k to N_eff_=99k (**Supplementary Figure 8**). We found, as expected, that increased sample size is associated with decreased estimated effect size distributions. Nevertheless, the estimates of discoverability for each sample size have overlapping 95% CIs and the standard error decreases with increasing sample size. Future brain structure GWAS in larger sample sizes may therefore lead to decreased discoverability with tighter estimates, but nevertheless we expect that those estimates will likely be contained within the CIs shown here, if the assumptions for the estimation procedure in GENESIS have been met. The sample sizes were relatively smaller in cortical/subcortical structures compared to behavioral traits (e.g. mean value of sample size is 32,512 for cortical surface area 193,640 for behavioral traits, *Welch’s t*-test p = 0.048) so future exploration of genetic architecture in increased sample sizes will help solidify the findings of decreased polygenicity and increased discoverability of brain structure traits relative to neuropsychiatric disorders and cognitive traits. Second, in this study we only tested two- or three-component mixture models to estimate polygenicity and discoverability, consistent with previous work [Zhang et al., 2018]. Given the inflations shown in QQ-plots for some traits (**Supplementary Figure 1-5**), more complex models (> 3 components) may better fit the effect size distributions. Finally, in this study we identified all sSNPs regardless of their genomic position or functional annotation. Future studies may explore discoverability within specific functional categories (e.g. enhancers present within a cell-type or context) to derive specific hypotheses about the mechanisms underlying trait variation [Johnson et al., 2020; Shadrin et al., 2020].

## Conclusion

Overall, our results use estimates of genetic architecture to test long-standing hypotheses about brain structure traits as endophenotypes for neuropsychiatric disorders.

## Acknowledgments

We express our gratitude to Drs. Nilanjan Chatterjee and Yan Zhang for their kind help in the use of GENESIS. JLS was supported by grants from the National Institute of Mental Health (NIMH) (R01MH118349, R01MH120125, R01MH121433). MIL was also supported by the NIMH(R01MH118349).

## Data Availability Statement

The data that support the findings of this study was derived from publicly available GWAS summary statistics described in the “*GWAS summary statistics”* in the Methods.

