## Supplementary Information for "Evaluating brain structure traits as endophenotypes using polygenicity and discoverability"

**Supplementary Figure** **2**

Supplementary Figure 1: Q-Q plots of model fit for cortical surface area 2
Supplementary Figure 2: Q-Q plots of model fit for cortical thickness 7
Supplementary Figure 3: Q-Q plots of model fit for subcortical volumes 12
Supplementary Figure 4: Q-Q plots of model fit for neuropsychiatric disorders, addiction relative traits and cognition 13
Supplementary Figure 5: Q-Q plots of model fit for anthropometric measurements 14
Supplementary Figure 6: Effect size distributions across cortical structures and subcortical volumes 15
Supplementary Figure 7: The relationship between discoverability, polygenicity and heritability 16
Supplementary Figure 8: Impacts on variance estimates by sample size 18

**Supplementary Table (header information)** **19**

Supplementary Table 1: Study summary 19
Supplementary Table 2: Estimated sSNPs and heritability from M2 model and model selection 19
Supplementary Table 3: Estimated sSNPs and heritability from M3 model 20
Supplementary Table 4: Predicted sample sizes required to explain the full heritability of traits 21

### Supplementary Figure

#### Supplementary Figure 1: Q-Q plots of model fit for cortical surface area

Upper panel (green) : M2, Lower panel (blue) : M3; “*” indicates best fit model
$\lambda_{obs}$ indicates the genomic control factor in the observed GWAS summary statistics
$\lambda_{fit}$ indicates the mean genomic control factor over 100 simulated datasets using estimated parameters from GENESIS

### Supplementary Figure 1: Q-Q plots of model fit for cortical surface area

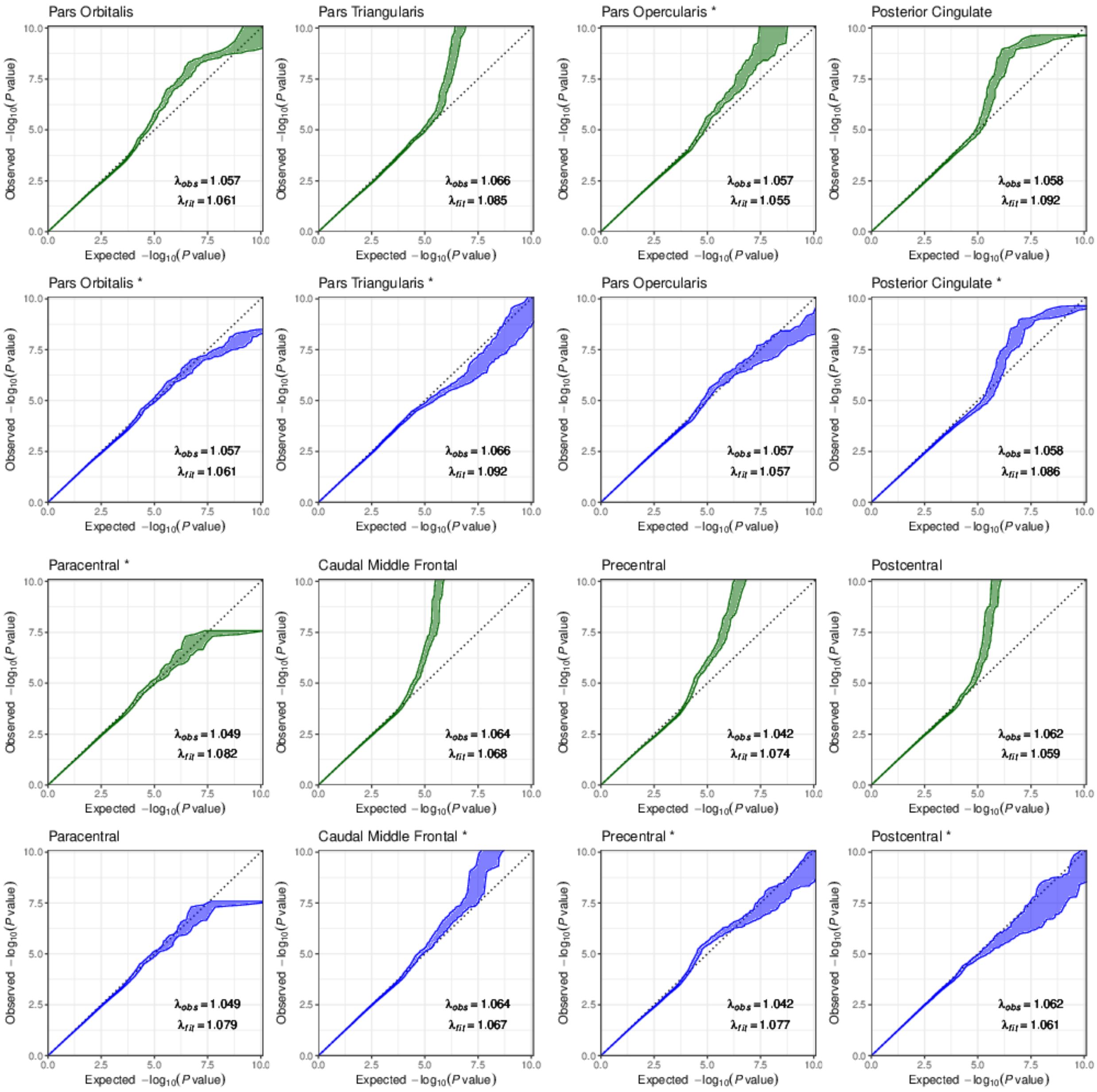

### Supplementary Figure 1: Q-Q plots of model fit for cortical surface area

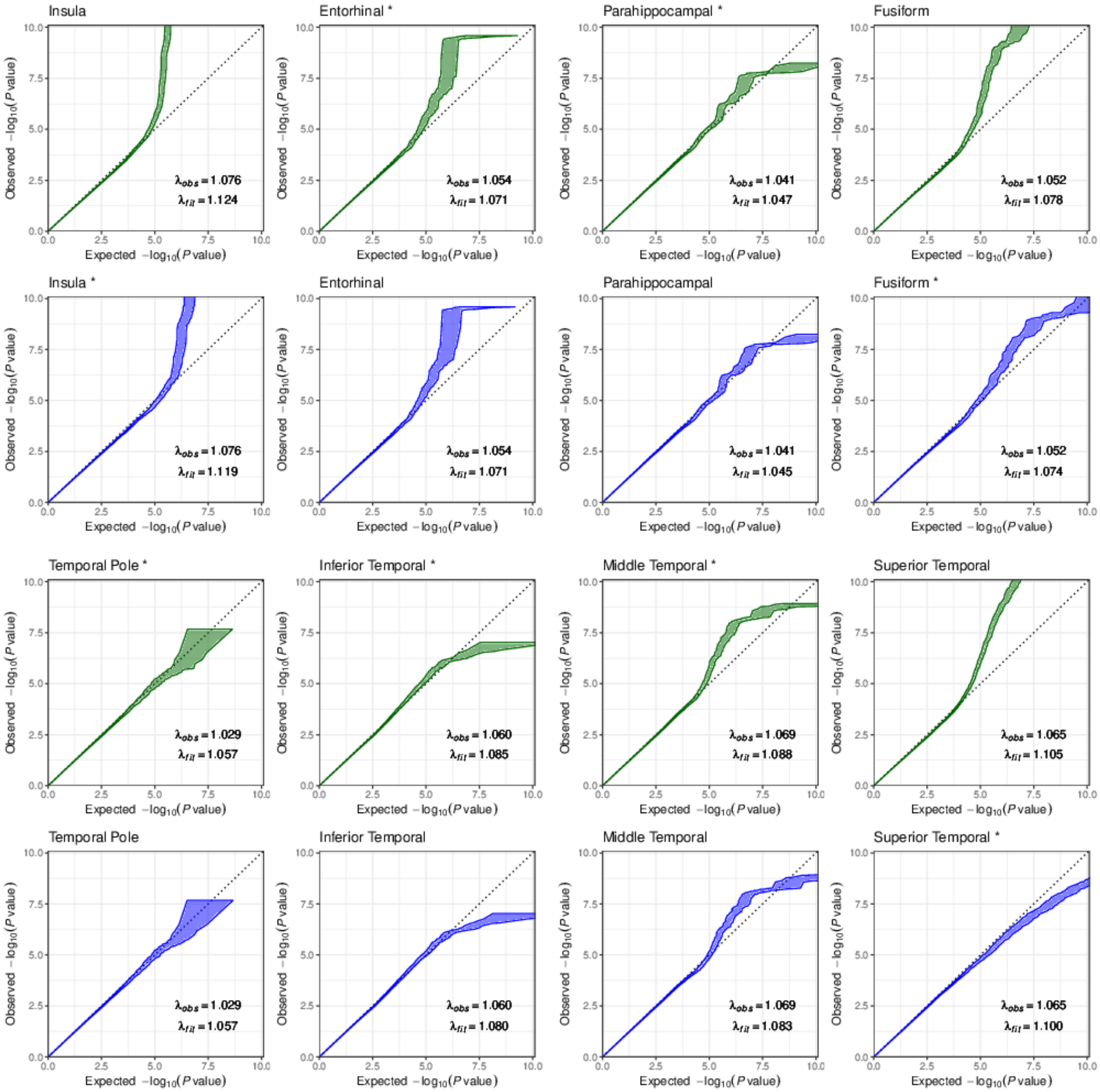

### Supplementary Figure 1: Q-Q plots of model fit for cortical surface area

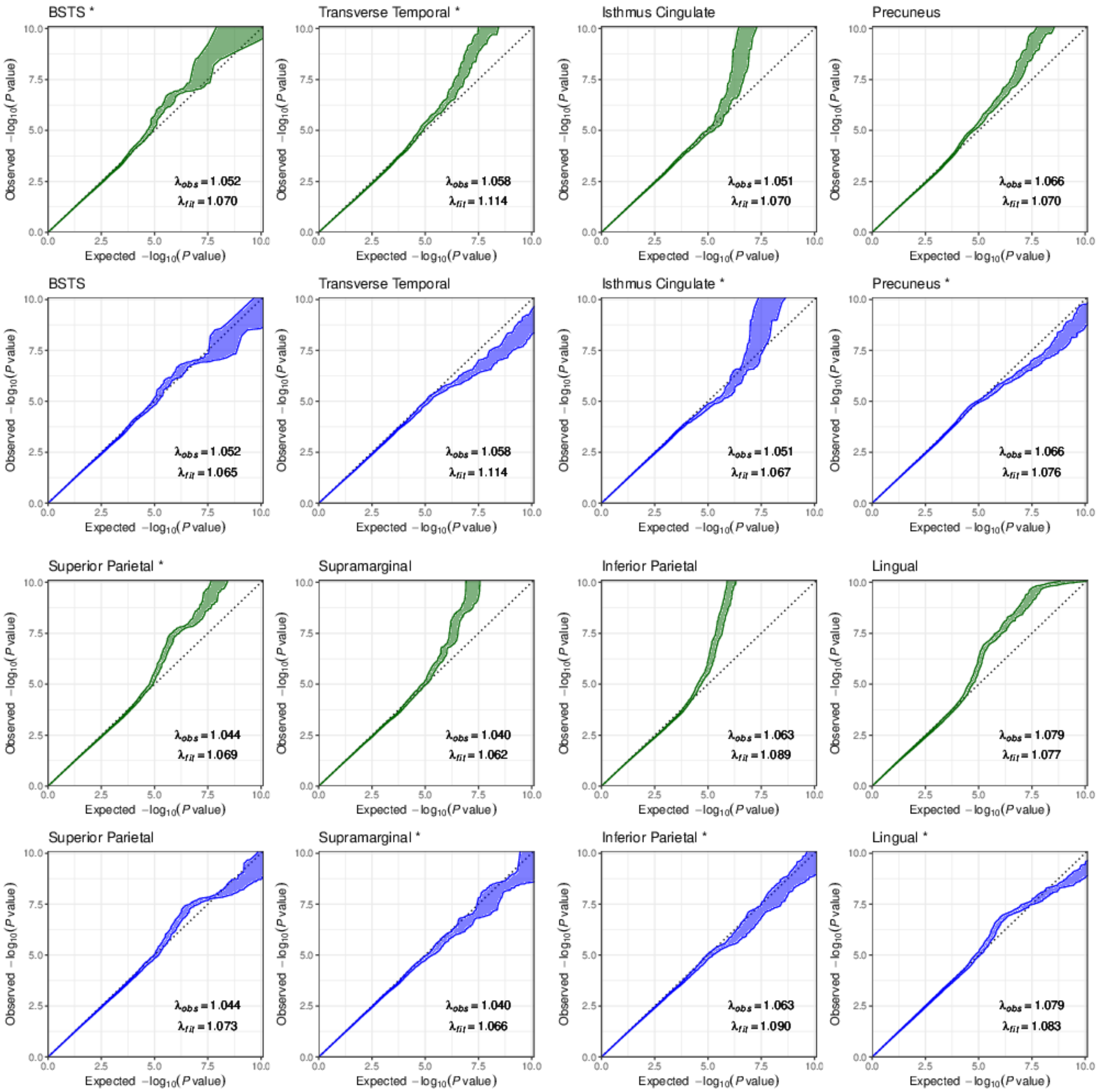

Supplementary Figure 1: Q-Q plots of model fit for cortical surface area

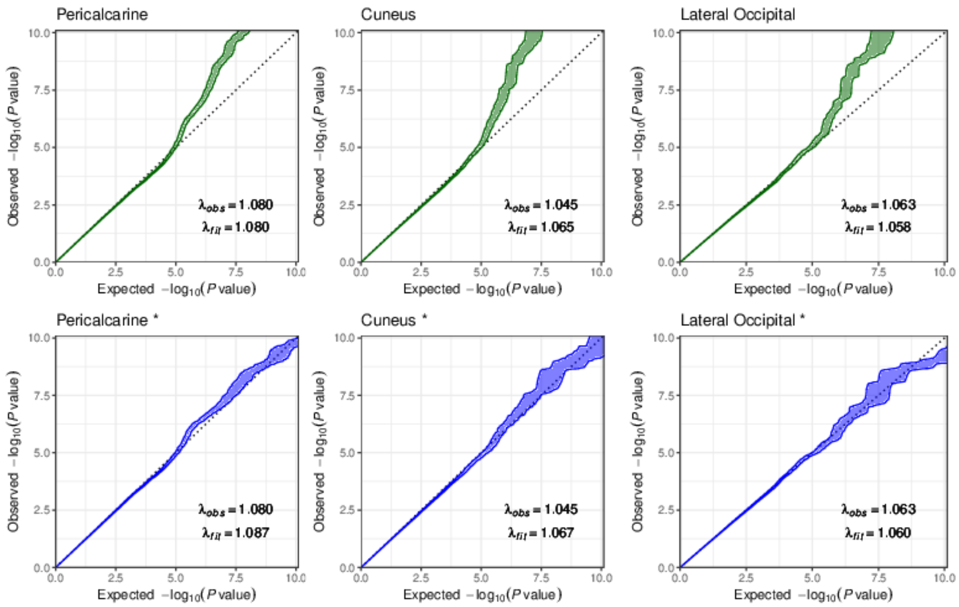

### Supplementary Figure 2: Q-Q plots of model fit for cortical thickness

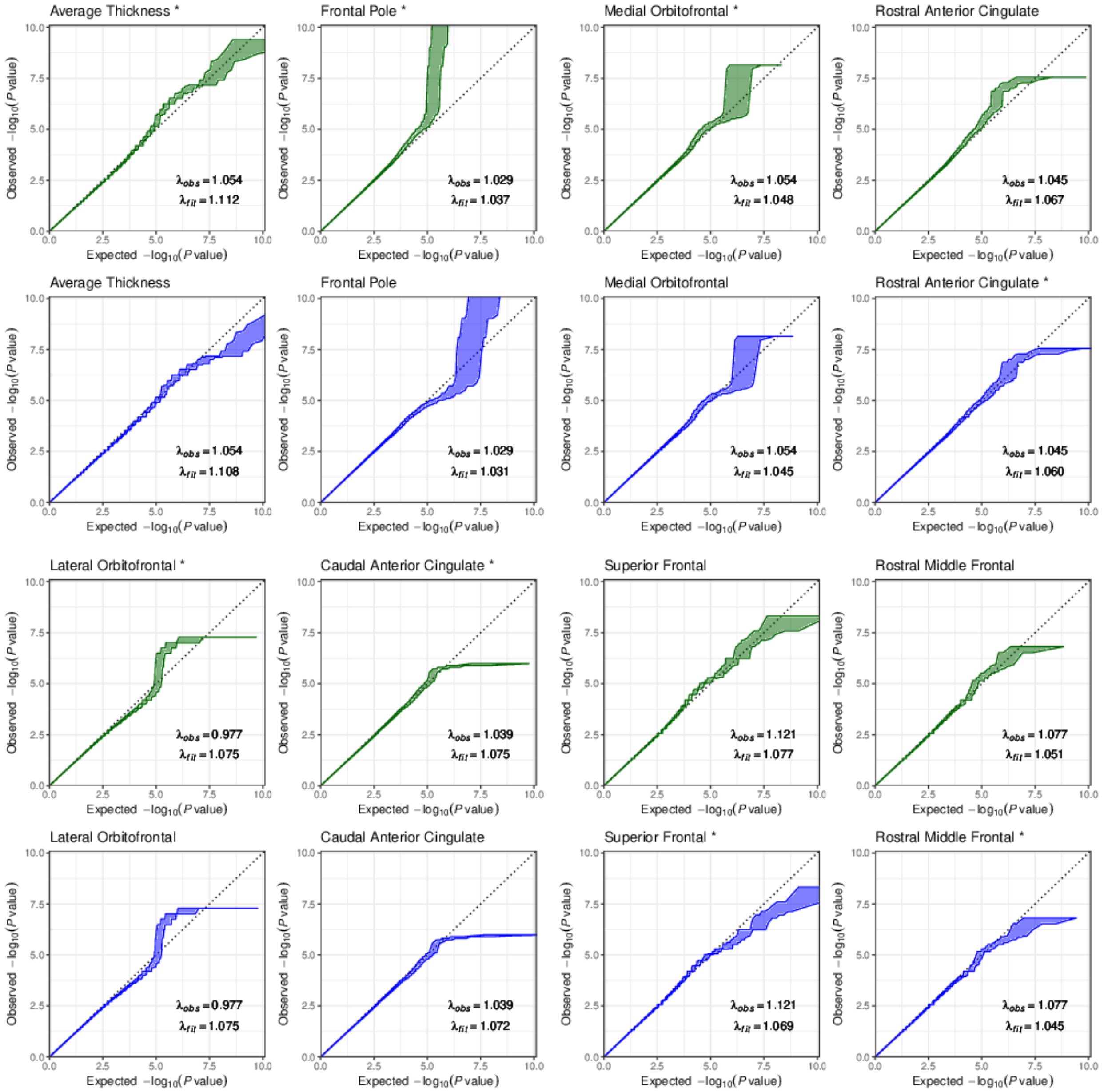

Supplementary Figure 2: Q-Q plots of model fit for cortical thickness

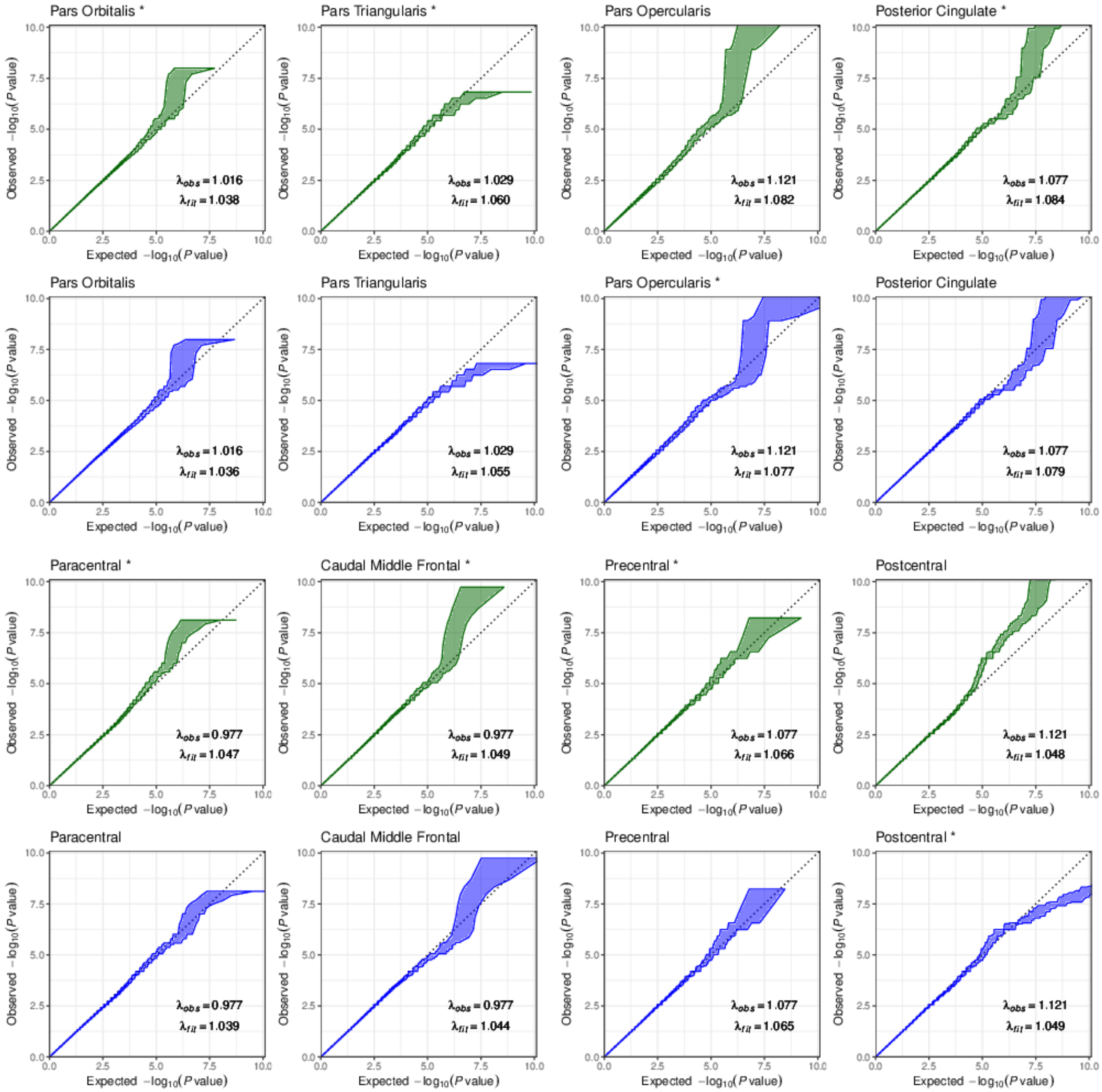

### Supplementary Figure 2: Q-Q plots of model fit for cortical thickness

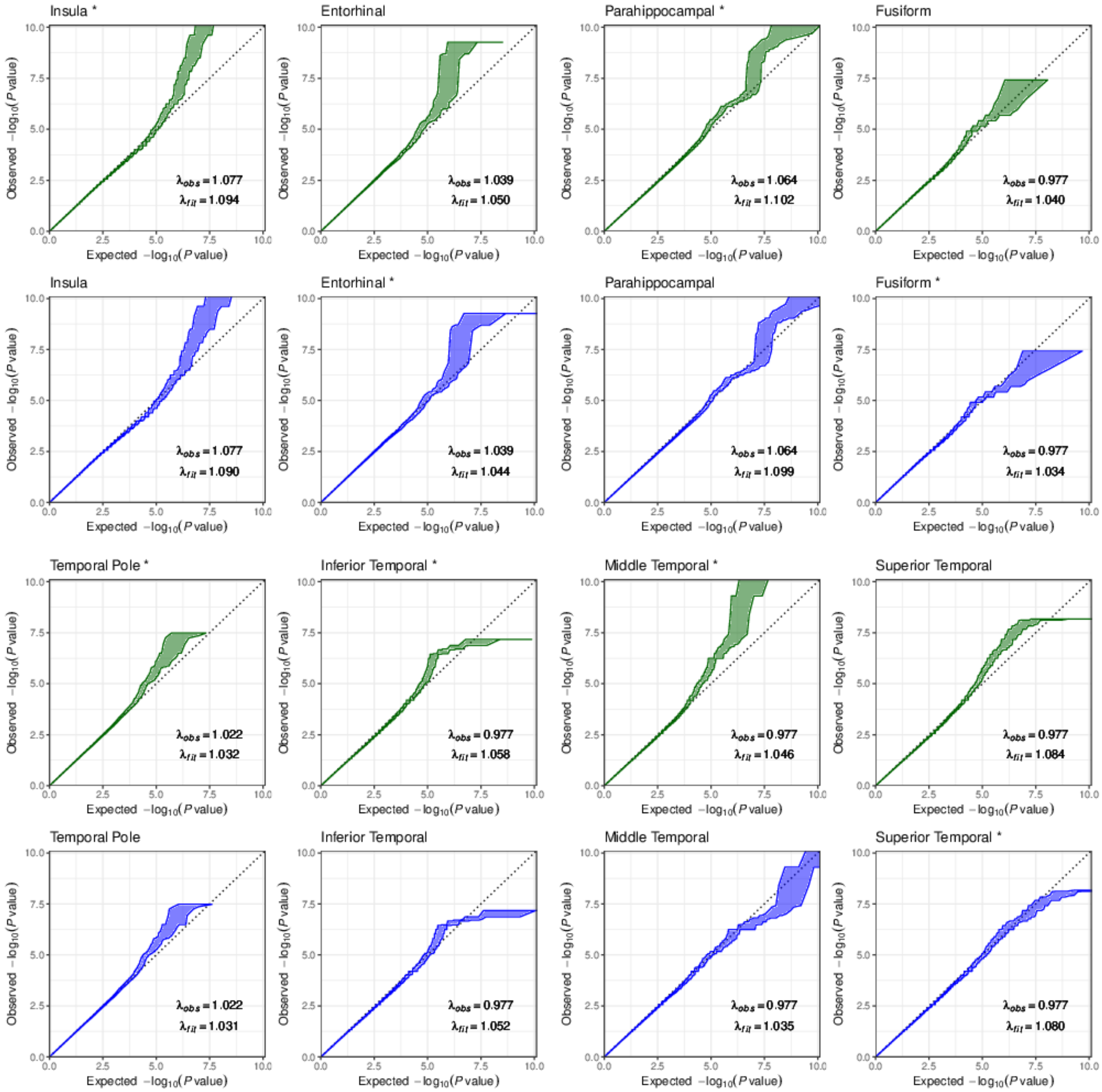

Supplementary Figure 2: Q-Q plots of model fit for cortical thickness

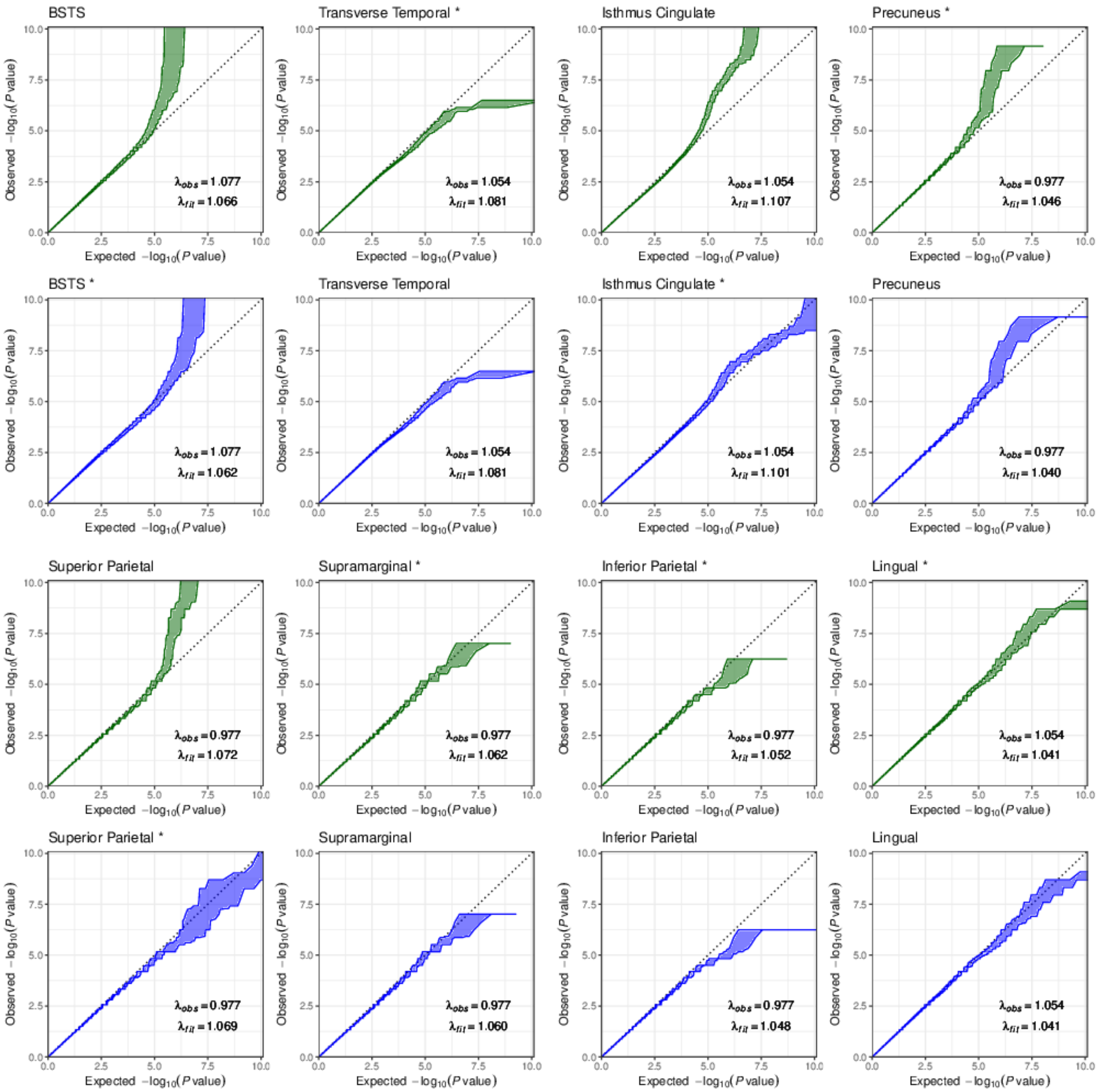

#### Supplementary Figure 2: Q-Q plots of model fit for cortical thickness

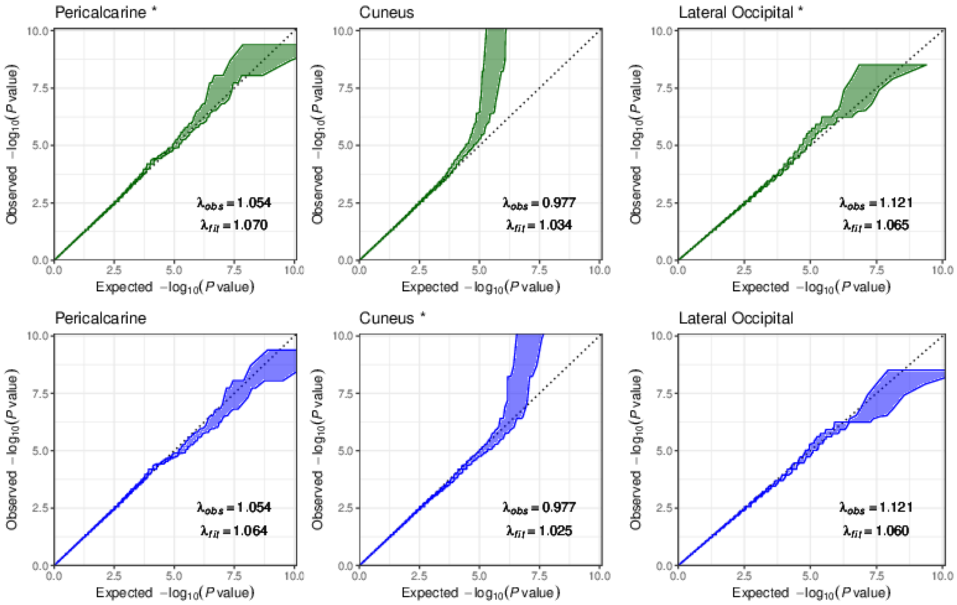

Supplementary Figure 3: Q-Q plots of model fit for subcortical volumes

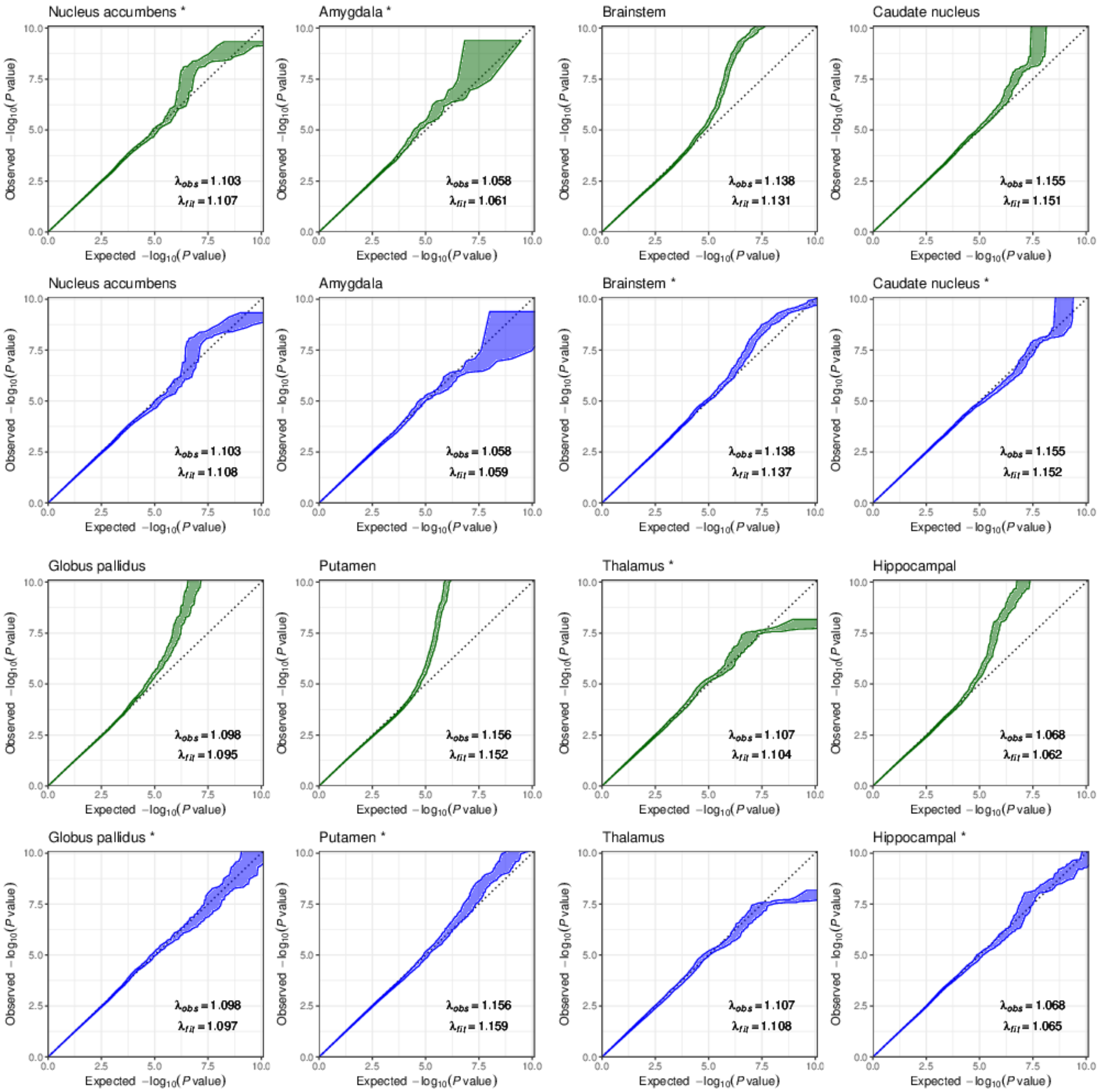

Supplementary Figure 4: Q-Q plots of model fit for neuropsychiatric disorders, addiction relative traits and cognition

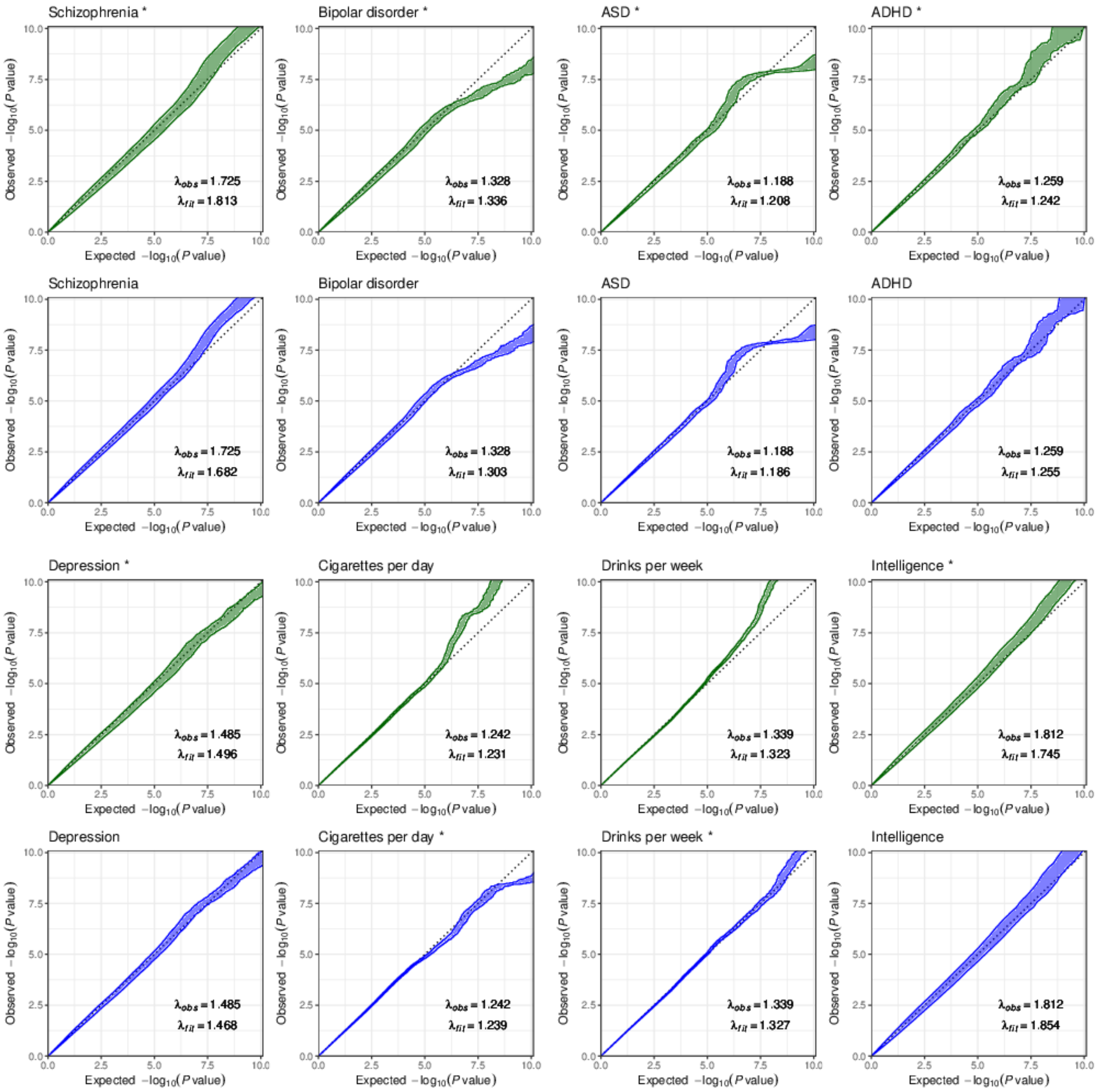

Supplementary Figure 4: Q-Q plots of model fit for neuropsychiatric disorders, addiction relative traits and cognition

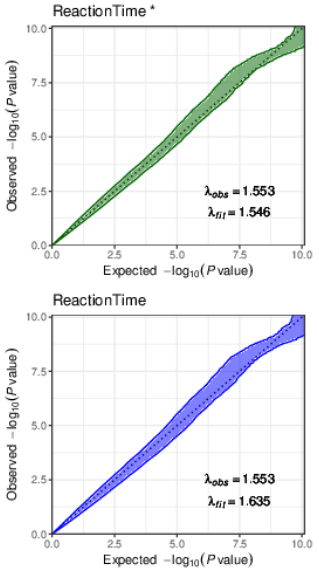


Supplementary Figure 5: Q-Q plots of model fit for anthropometric measurements

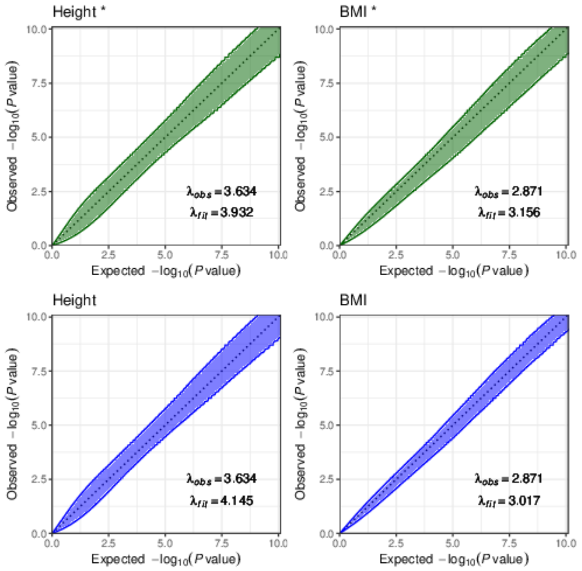

### Supplementary Figure 6: Effect size distributions across cortical structures and subcortical volumes

(**a**) Increased effect size in cortical surface area compared to cortical thickness. In the forest plots, the 50th percentile of ranked sSNP absolute effect size is shown with 95% CIs as error bars. * indicates phenotypes with lower/upper limits of proportion of sSNPs in cluster 1 out of range (>1 or <0, limited to 1 or 0). $ indicates phenotype where the 95% CI lower limit of $\sigma_1^2$ or $\sigma_2^2$ had a negative value and was limited to 0. Thus, the CI for those phenotypes needs caution in interpretation. (**b**) The point estimates in (**a**) were mapped to the cortical regions. (**c**) Effect size distribution comparing across cortical structures and subcortical volumes.

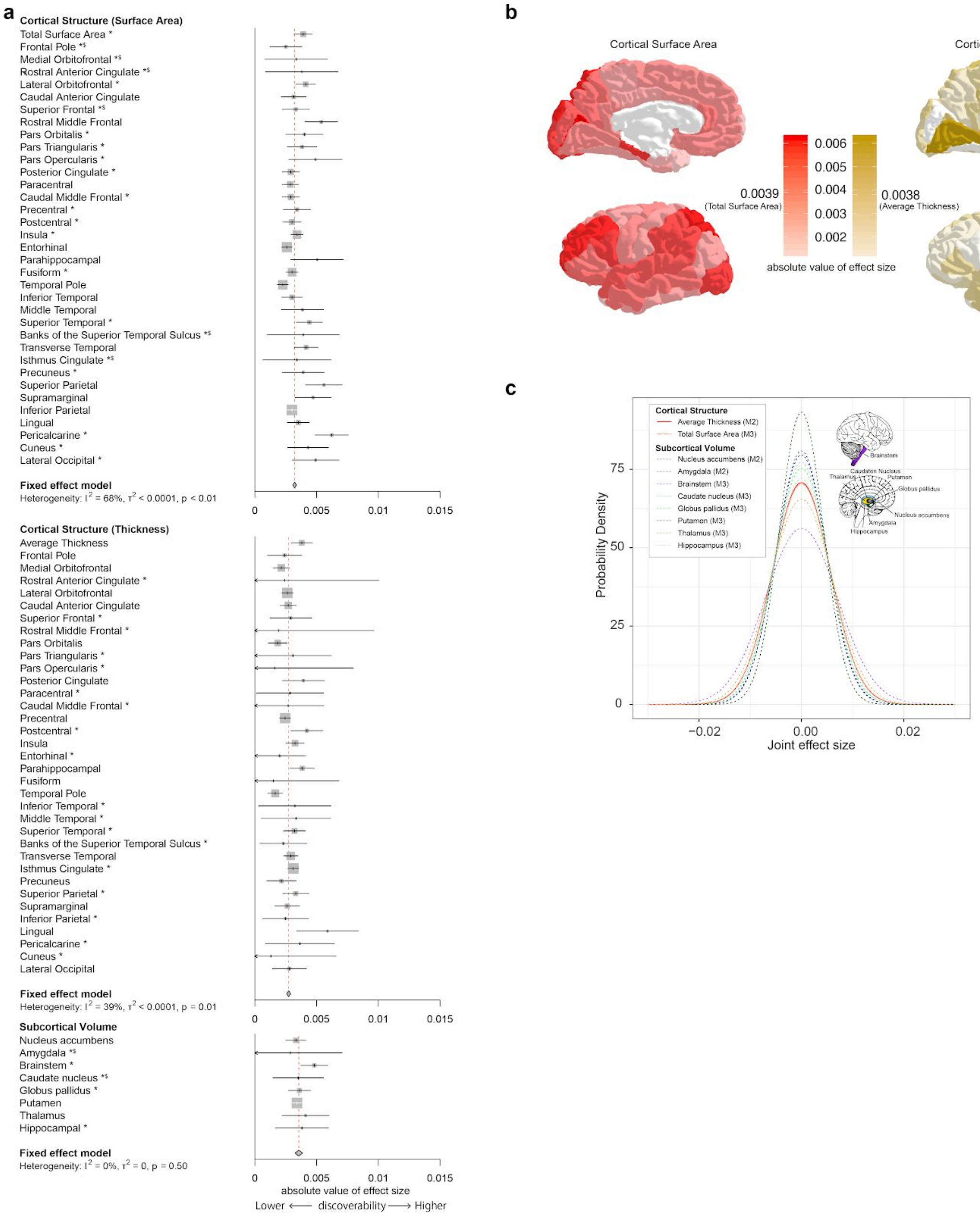

### Supplementary Figure 7: The relationship between discoverability, polygenicity and heritability

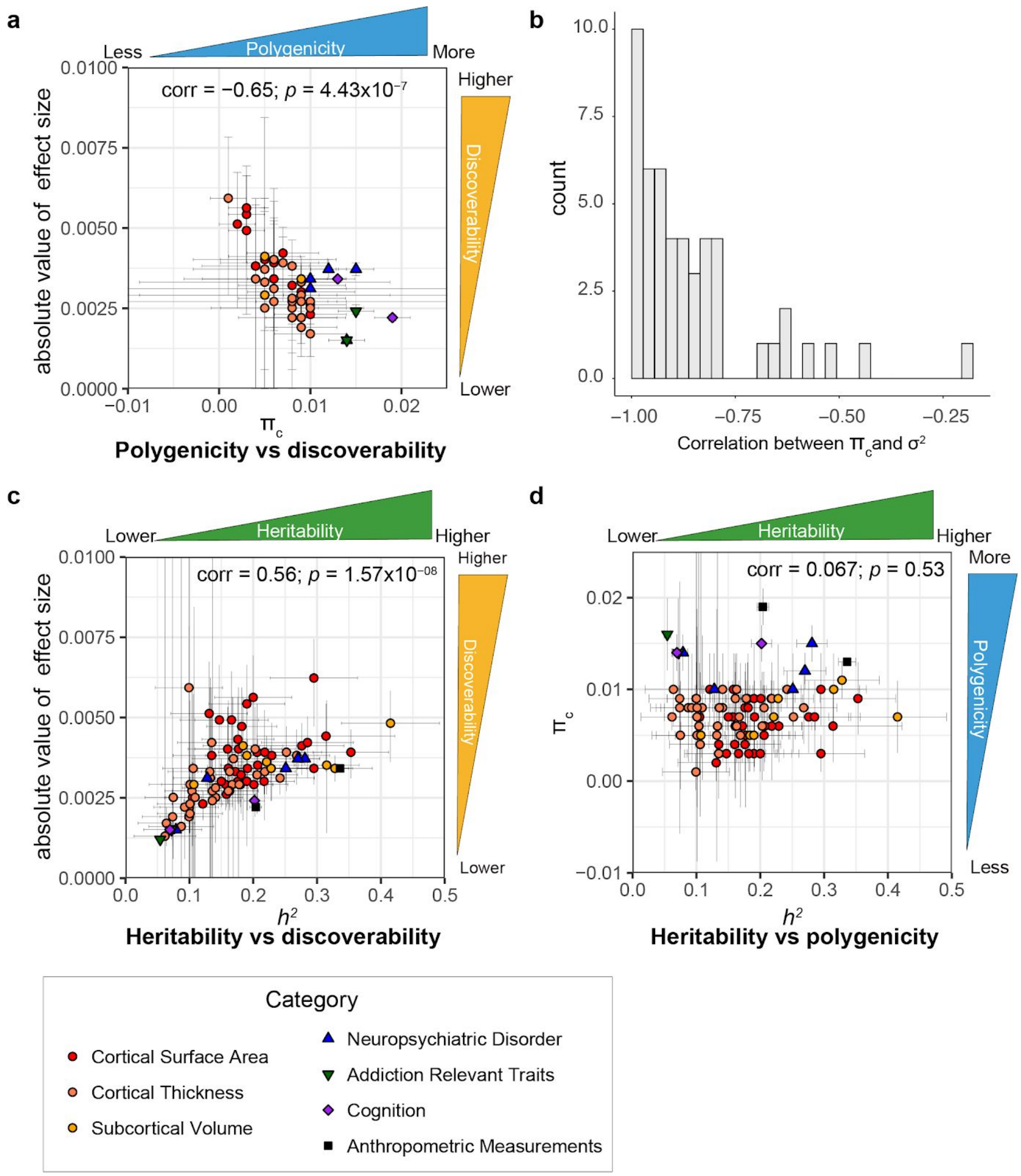


Pearson's correlation coefficient (corr) showed (**a**) significant negative correlation between polygenicity ($\pi_c$) and discoverability (absolute value of effect size). (**b**) Covariance matrix from GENESIS output indicated a negative correlation between estimates of $\pi_c$ and $\sigma^2$ which is likely producing the negative correlation in (**a**). We also observed (**c**) significant positive correlation between heritability ($h^2$) and discoverability, but (**d**) no significant correlation between heritability and polygenicity. Only phenotypes best fit to M2 were shown in (**a**) to simplify assessment of the correlation of estimated parameters in the model **(b)**. Error bar indicates 95% CIs of the estimate.

#### Supplementary Figure 8: Impacts on variance estimates by sample size

Three schizophrenia GWASs (Ripke et al. 2013; Ripke et al. 2014; Pardiñas et al. 2018) with different sample sizes ($N_{eff}$ = 31,519 ~ 99,863) were compared. Effect sizes were decreased with larger sample numbers (**a-c**). Effective sample size was calculated by 4/(1/cases+1/controls) (Willer et al. 2010).

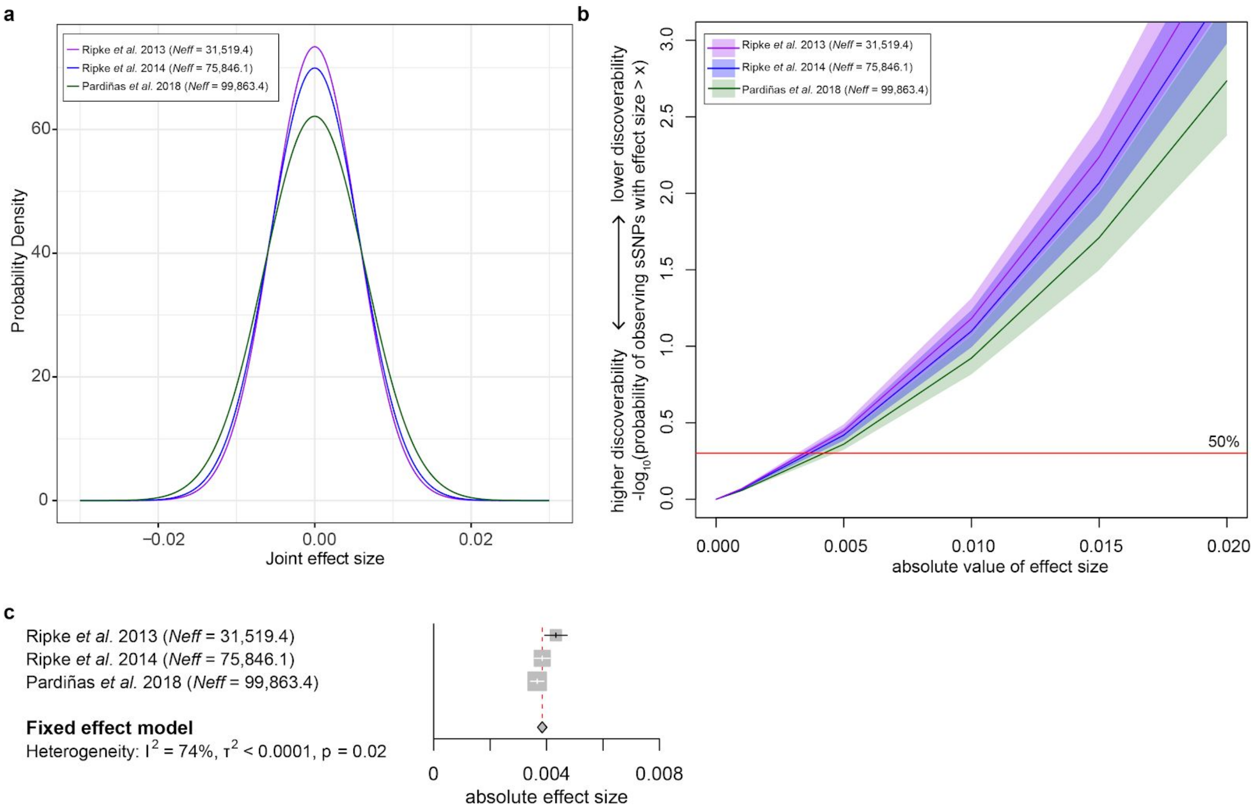

### Supplementary Table (header information)

#### Supplementary Table 1: Study summary

| Header name | Description |
|---|---|
| Category | category (e.g. Cortex (Surface Area)) |
| Trait | trait (e.g. Total Surface Area) |
| Case # | number of case samples (within case vs control study only) |
| Control # | number of control samples (within case vs control study only) |
| Total # (effective #) | total sample size (for case vs control study, effective sample number was calculated by 4/(1/case+1/control) |
| Ref | Reference |

#### Supplementary Table 2: Estimated sSNPs and heritability from M2 model and model selection

| Header name | Description |
|---|---|
| Category | category (e.g. Cortex (Surface Area)) |
| Trait | trait (e.g. Total Surface Area) |
| GWAS Marker | # of GWAS SNPs after QC |
| # of sSNPs | number of susceptibility SNPs (and standard error) |
| pi_c | proportion of sSNPs (and standard error) |
| Heritability | heritability estimates (and standard error) |
| BIC_M2 | modified BIC based on fit of the2-component model (M2) |
| BIC_M3 | modified BIC based on fit of the 3-component model (M3) |
| Ratio | ratio of two variance estimates from M3 |
| Best-fit Model | final choice of best fit model (see Methods) |

Supplementary Table 3: Estimated sSNPs and heritability from M3 model

| Header name | Description |
|---|---|
| Category | category (e.g. Cortex (Surface Area)) |
| Trait | trait (e.g. Total Surface Area) |
| GWASMaker | # of GWAS SNPs after QC |
| # of sSNPs | number of susceptibility SNPs (and standard error) |
| pi_c | proportion of sSNPs (and standard error) |
| Proportion of sSNPs in cluster 1 | proportion of sSNPs (and standard error) in cluster 1 (larger effect sizes) |
| Heritability in cluster 1 | heritability estimates (and standard error) explained by sSNPs in cluster 1 |
| Heritability in cluster 2 | heritability estimates (and standard error) explained by sSNPs in cluster 2 |
| Total Heritability | heritability estimates (and standard error) explained by all sSNPs |

#### Supplementary Table 4: Predicted sample sizes required to explain the full heritability of traits

| Header name | Description |
|---|---|
| Category | category (e.g. Cortex (Surface Area)) |
| Trait | trait (e.g. Total Surface Area) |
| model | best fit model (M2 or M3) |
| Required sample # | # of GWAS sample required to pass 99% of heritability explained by GWS sSNPs |
| % of GV | % of heritability explained by GWS sSNPs at 20 M individuals when it is not expected to pass 99% (in Required sample # column) |
